# Use of Therapeutic Drug Monitoring to Characterize Cefepime-Induced Neurotoxicity

**DOI:** 10.1101/2020.08.13.250456

**Authors:** Veena Venugopalan, Cara Nys, Natalie Hurst, Yiqing Chen, Maria Bruzzone, Kartikeya Cherabuddi, Nicole Iovine, Jiajun Liu, Mohammad H. Al-Shaer, Marc H. Scheetz, Nathaniel Rhodes, Charles A. Peloquin, Kenneth Klinker

## Abstract

**Background:** The incidence of cefepime-induced neurotoxicity (CIN) in hospitalized patients is highly variable. Although greater cefepime exposures incite neurotoxicity, data evaluating trough thresholds associated with CIN remains limited. The objectives of this study were to evaluate the incidence of CIN, assess the relationship between cefepime trough concentrations and CIN, investigate clinical factors associated with CIN, and describe electroencephalogram (EEG) abnormalities in CIN.

**Methods:** This was a retrospective study of adult patients who had received ≥ 5 days of cefepime with ≥ 1 trough concentration > 25 mg/L. Potential CIN cases were identified utilizing neurological symptoms, neurologist assessments, EEG findings and improvement of neurotoxicity after cefepime discontinuation.

**Results:** One-hundred and forty-two patients were included. The incidence of CIN was 13% (18/142). The mean cefepime trough concentration in CIN patients was significantly greater than the non-neurotoxicity group (74.2 mg/L ± 41.1 vs. 46.6 mg/L ± 23, p=0.015). Lower renal function (creatinine clearance < 30 ml/min), greater time to therapeutic drug monitoring (TDM) (≥72 hours), and each 1 mg/mL rise in cefepime trough were independently associated with increased risk of CIN. Moderate generalized slowing of the background rhythm was the most common EEG pattern associated with CIN.

**Conclusion:** Cefepime should be used cautiously in hospitalized patients due to the risk of neurotoxicity. Patients with greater renal function and those who had early cefepime TDM (≤ 72 hours) had lower risk of CIN.

## Introduction

Cefepime-induced neurotoxicity (CIN) is a well-documented adverse effect.^1^ A systematic review reported an incidence of 15% in the intensive care unit (ICU) population whereas rates as high as 23% have been reported in hospitalized adults.^2,3^ In 2012, the Food and Drug Administration issued a safety communication warning of the risk of non-convulsive status epilepticus with cefepime use, particularly in patients with renal impairment.^4^ The pathophysiology of cefepime neurotoxicity is thought to be related to the concentrationdependent inhibition at GABA-A receptors, resulting in central excitotoxicity.^5,6^ The most common clinical features associated with CIN are diminished level of consciousness, disorientation or agitation, and myoclonus.^5^

Although widely accepted that an increase in cefepime exposure potentiates CIN, the exact relationship between cefepime concentration and neurotoxicity has not been fully elucidated. The objectives of this study were to evaluate the incidence CIN in our patient population, assess the relationship between cefepime trough concentrations and CIN, investigate clinical factors associated with development of neurotoxicity, and describe electroencephalogram (EEG) abnormalities in CIN.

## Materials and Methods

### Study Design

This was a retrospective, cohort study from March 2016 to May 2018 at the University of Florida Health Shands Hospital, which is a 1,162-bed tertiary academic medical center in Gainesville, Florida. Beta-lactam TDM has been available at our institution since 2016. Testing however, is not routinely performed on all patients but is obtained at the discretion of the treating physician and pharmacist. The study investigators wanted to evaluate CIN in a high-risk group. Thus, this study included adult patients who had received ≥ 5 days of cefepime therapy, and had at least one cefepime trough concentration > 25 mg/L. Based on prior studies indicating increased probability of CIN when free trough concentrations were >22 mg/ml, only those patients with cefepime troughs above 25 mg/L were included.^7,8^ Two Infectious Diseases specialists (KC, NI) and an Infectious Diseases pharmacist (VV), independently reviewed medical records for patients with suspected CIN to determine if they met the case definition for the study. EEGs were performed based on the International Federation of Clinical Neurophysiology (IFCN) guidelines and results were interpreted by a board certified epileptologist (MB).

Cefepime plasma concentrations were obtained at steady state (≥ 24 hours from initiation of cefepime). For each sample, an aliquot of 4 ml of blood was drawn into non-heparinized tubes, which were centrifuged at 1000g for 10 minutes, and the resulting plasma stored at −80C. Plasma concentrations of cefepime were measured in the Infectious Diseases Pharmacokinetics Lab (IDPL) at the University of Florida, using a validated ultrahigh pressure liquid chromatography assay with triple quadrupole mass spectroscopy (LC-MS-MS).^9^

Patient demographics, comorbid conditions, laboratory values, cefepime dosing, drug concentrations, and in-hospital mortality were extracted from the electronic medical records, Epic® 2015 software (Verona, Wisconsin). The study was approved by the Institutional Review Board at the University of Florida.

### Outcomes and Definitions

The primary outcomes of the study were to quantify the incidence of CIN in our patient population based on pre-defined criteria and to assess the association between cefepime trough concentrations and neurotoxicity. The secondary outcomes of the study were to investigate the clinical factors associated with development of CIN and to determine if a correlation exists between CIN and in-hospital mortality.

To be considered a potential CIN case, patients were required to fulfill ≥ 2 of the National Cancer Institute (NCI) criteria for neurological toxicity.^10^ The NCI criteria includes symptoms such as presence of new onset confusion, delirium, and drowsiness. CIN was defined as a patient meeting two of the following three criteria: 1) neurology consult describing CIN, 2) electroencephalogram (EEG) findings consistent with cefepime toxicity, 3) and improvement of signs and symptoms of neurotoxicity after cefepime discontinuation. The occurrence of all potentially drug-related adverse events were systematically assessed.

The observed EEG findings were categorized based on the 2012 American Clinical Neurophysiology Society standardized EEG critical care terminology.^11^ EEG abnormalities were classified as follows: lateralized periodic discharges (LPDs), generalized periodic discharges (GPDs) with and without triphasic morphology, generalized rhythmic delta activity (GRDA), and multifocal sharps and spike-and-waves (MfSWs). Non-convulsive status epilepticus was defined electrographically as epileptiform discharges (ED) at >2.5Hz or EDs ≤ 2.5 Hz or rhythmic delta/theta activity (>0.5 Hz) AND one of the following: EEG and clinical improvement after intravenous antiepileptic drugs (AEDs), or subtle clinical ictal phenomena during the EEG patterns mentioned above, or typical spatiotemporal evolution, without associated clinical convulsions.^12^

### Statistical Analysis

Comparisons between the non-NT and CIN group was performed using independent Student’s t-test for continuous data with normal distribution and Chi-square test or Fisher’s exact test for categorical data as appropriate; the Mann-Whitney test was used to compare medians for continuous variable not normally distributed. To evaluate cefepime concentrations between the non-NT and CIN groups, a generalized linear model was conducted with prespecified covariates including age, ICU at time of concentration, length of hospital stay, serum creatinine, duration of cefepime therapy, and total daily dose of cefepime. Unadjusted and adjusted binary logistic regression models were conducted using neurotoxicity as the dependent variable and age, sex, renal function, ICU status, cefepime trough concentration, and time to TDM as independent variables. P-values <0.05 were considered statistically significant. Statistical analysis was performed on JMP Pro v15.0 (SAS Institute, Cary, NC).

## Results

One-hundred and forty-two patients were included in the analysis. Baseline patient characteristics are summarized in Table 1. The incidence of CIN was 13% (18/142) among patients with trough concentrations > 25 mg/L. The mean patient age in the CIN group was 62.9 ± 15.8 years. Hypertension as a comorbid condition was more frequent in the CIN group (88.9%) vs. the non-neurotoxic (non-NT) group (61.3%) (p=0.03). Markers of renal function (serum creatinine and creatinine clearance) were obtained at the time of therapeutic drug monitoring. All neurotoxic patients in the CIN group (n=18) had acute or chronic kidney function. Patients exhibiting cefepime neurotoxicity had lower creatinine clearance compared to those who did not (38 ± 18.4 ml/min vs. 82.1 ± 44.0 ml/min, p<0.0001). Sixty-one percent (n=11) with CIN were in an ICU setting at the time that the cefepime concentration was obtained. Hospital and ICU length of stay (LOS) did not vary between non-NT and CIN groups.

**Table 1.** Patient demographics and Baseline Characteristics

| Characteristics | Overall<br>(n=142) | Non-NT<br>(n=124) | CIN<br>(n=18) | p-value |
| --- | --- | --- | --- | --- |
| Age, years | 57.8 ± 16.6 | 57.1 ± 16.7 | 62.9 ± 15.8 | 0.17 |
| Male | 88 (62) | 77 (62) | 11 (61) | 1 |
| BMI<sub>a</sub>, kg/m<sup>2</sup> | 29.9 ± 9.8 | 29.9 ± 10 | 29.8 ± 8.4 | 0.95 |
| Comorbid conditions |  |  |  |  |
| Hypertension | 92 (65) | 76 (61) | 16 (89) | 0.03 |
| Diabetes mellitus | 64 (45) | 53 (43) | 11 (61) | 0.20 |
| Renal dysfunction<sub>b</sub> | 82 (58) | 64 (52) | 18 (100) | <0.0001 |
| AKI<sub>c</sub> | 66 (47) | 51 (41) | 15 (83) | 0.0009 |
| CKD<sub>d</sub> | 57 (40) | 45 (36) | 12 (67) | 0.02 |
| CVA<sub>e</sub> | 13 (9) | 10 (8) | 3 (17) | 0.22 |
| Seizure disorder | 18 (13) | 16 (13) | 2 (11) | 1 |
| Hemodialysis | 18 (13) | 14 (11) | 4 (22) | 0.25 |
| CrCl, ml/min<sub>f</sub> | 76.5 ± 44.1 | 82.1 ± 44.0 | 38 ± 18.4 | <0.0001 |
| SCr (all patients)<sub>g</sub> | 1.4 ± 1.1 | 1.3 ± 1 | 2.3 ± 1.5 | <0.0001 |
| SCr (non-dialysis)<sub>g</sub> | 1.3 ± 1 | 1.2 ± 0.9 | 2.0 ± 1.2 | 0.0004 |
| Mechanical ventilation | 68 (48) | 62 (51) | 6 (33) | 0.21 |
| Concomitant medications |  |  |  |  |
| Anticonvulsants | 37 (26) | 34 (27) | 3 (17) | 0.40 |
| Benzodiazepines | 34 (24) | 31 (25) | 3 (17) | 0.56 |
| Other antibiotics | 114 (80) | 99 (80) | 15 (83) | 1 |
| LOS<sub>h</sub> prior to TDM<sub>i</sub>, days | 15.2 ± 16.3 | 15.9 ± 17.1 | 10.5 ± 6.5 | 0.36 |
| ICU<sub>j</sub> at time of TDM<sub>i</sub> | 97 (68) | 86 (69) | 11 (61) | 0.59 |
| EEG | 31 (22) | 19 (15) | 12 (67) | <0.0001 |

\*Data are presented as mean (STD) or n (%); <sub>a</sub>BMI: Body mass index; <sub>b</sub>Renal dysfunction = AKI and/or CKD; <sub>c</sub>AKI: Acute kidney injury; <sub>d</sub>CKD: Chronic kidney disease; <sub>e</sub>CVA: cerebrovascular accident; <sub>f</sub>CrCl: Creatinine clearance; <sub>g</sub>SCr: Serum creatinine; <sub>h</sub>LOS: Length of stay; <sub>i</sub>TDM: Therapeutic drug monitoring; <sub>j</sub>ICU: Intensive care unit

Mean daily cefepime doses were significantly greater in the non-NT arm compared to those who experienced CIN (4.9 g/day ± 1.5 vs. 3.9 g/day ± 1.2 g/day, p=0.007) (Table 2). Prolonged infusion strategies (180- or 240-minute infusions) were more commonly utilized in the non-NT vs. CIN group (37% vs. 6%, p=0.0067). It was observed that patients with CIN had shorter mean duration of cefepime treatment relative to those with non-NT, however this was a non-significant difference (10.7 days ± 5.3 vs. 13.4 days ± 10.1, p=0.09). The mean time to cefepime neurotoxicity from the start of cefepime therapy was 7.4 ± 3.9 days. Measurements of cefepime concentrations were performed on average 6.5 days after the start of cefepime therapy. In 14 patients (78%) of CIN cases (data not shown), cefepime therapy was stopped within 72 hours of the onset of neurotoxic symptoms.

**Table 2.** Cefepime dosing, duration, and infusion strategies

| Characteristic | Overall<br>(n=142) | Non-NT<br>(=124) | CIN<br>(n=18) | p-value |
| --- | --- | --- | --- | --- |
| Cefepime daily dose, g/day | 4.8 ± 1.5 | 4.9 ± 1.5 | 3.9 ± 1.2 | 0.007 |
| Infusion strategy |  |  |  | 0.0067 |
| Intermittent± | 96 (67) | 78 (63) | 17 (94) |  |
| Extended# | 47 (33) | 46 (37) | 1 (6) |  |
| Duration of treatment, days | 13.1 ± 9.7 | 13.4 ± 10.1 | 10.7 ± 5.3 | 0.09 |
| Time to cefepime neurotoxicity from start of therapy, days | - | - | 7.4 ± 3.9 | - |
| Time from cefepime start to TDM, days | 5.6 ± 4.2 | 5.4 ± 4.4 | 6.5 ± 2.7 | 0.17 |

\*Data are presented as mean (STD) or n (%); TDM = therapeutic drug monitoring ±30-minute infusion; #180 or 240-minute infusion

Evaluating the primary outcomes of the study, the mean cefepime trough concentrations in those with CIN were significantly greater than those without neurotoxicity (74.2 mg/L ± 41.1 vs. 46.6 mg/L ± 23, p=0.015) (Figure 1, Table 3). Although not statistically significant, there was a trend towards greater in-hospital mortality in the CIN group (33%) compared with the non-NT group (15%) (Table 3). Lower renal function (CrCl < 30 ml/min), greater time to TDM (≥72 hours), and each 1 mg/mL incremental rise in cefepime trough were independently associated with increased risk of CIN in the adjusted regression analysis (Table 4).

**Figure 1.**
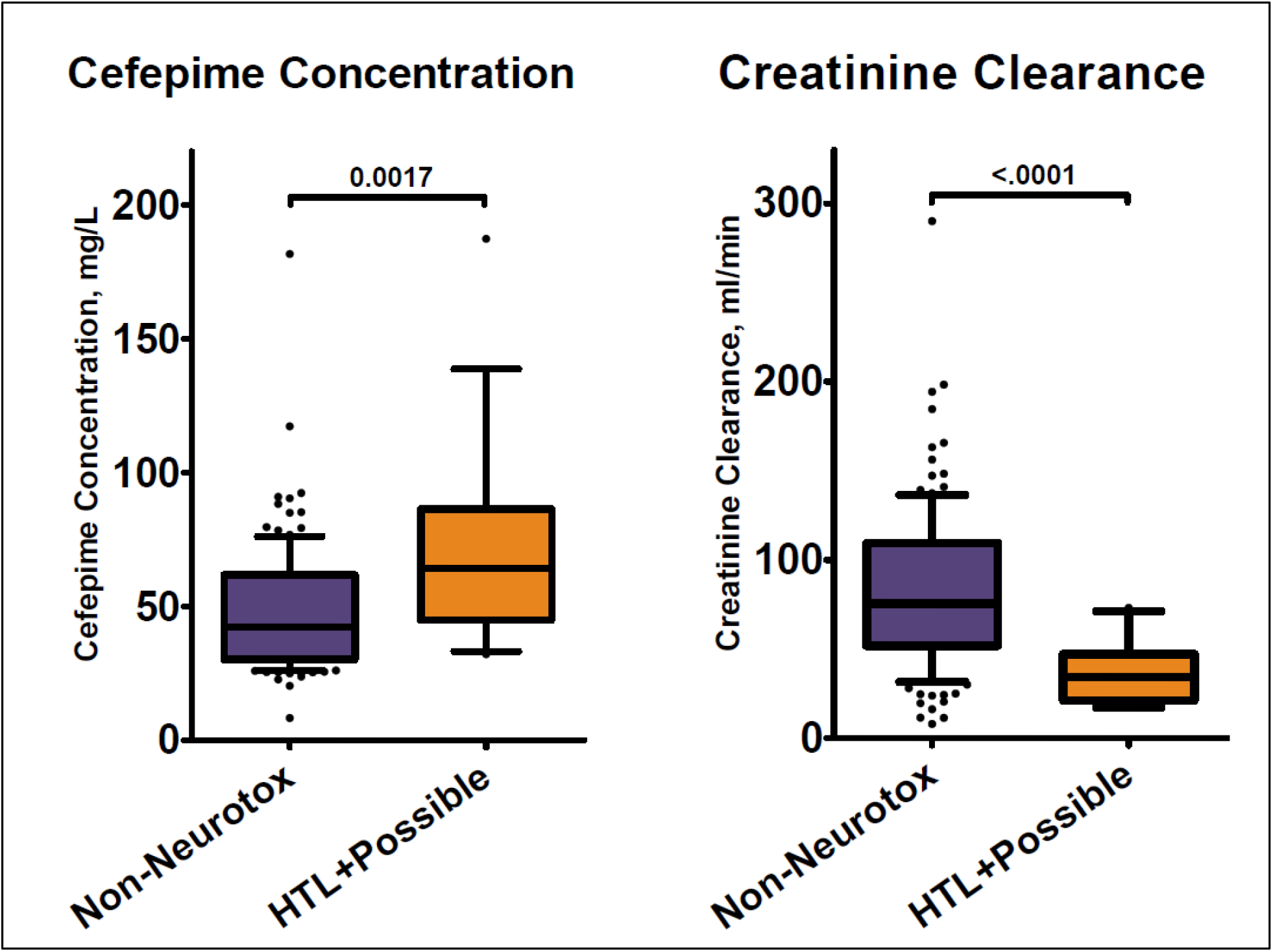

**Table 3.** Primary and secondary outcomes

| Characteristics | Overall<br>(n=142) | Non-NT<br>(n=124) | CIN<br>(n=18) | p-value |
| --- | --- | --- | --- | --- |
| Cefepime trough,<br>mg/L, mean ± SD | 51 ± 27.3 | 46.6 ± 23 | 74.2 ± 41.1 | 0.015 |
| In-hospital<br>mortality, n (%) | 24 (17) | 18 (15) | 6 (33) | 0.084 |
| CIN = cefepime-induced neurotoxicity; SD = standard deviation |  |  |  |  |

**Table 4.** Logistic regression analysis of the development of cefepime-induced neurotoxicity based on clinical characteristics

|  |  |  | Unadjusted analysis |  |  | Adjusted analysis |
| --- | --- | --- | --- | --- | --- | --- |
| Characteristic | OR | 95% CI | p- value | OR | 95% CI | p- value |
| Age (≥ 60 years) | 1.29 | 0.48-3.72 | 0.62 | 1.28 | 0.40-4.37 | 0.68 |
| Sex (Male) | 0.96 | 0.35-2.77 | 0.94 | 1.32 | 0.39-4.92 | 0.66 |
| CrCl (≥ 30 ml/min) | 0.12 | 0.04-0.37 | 0.0004 | 0.15 | 0.04-0.58 | 0.0063 |
| Time from cefepime start to TDM (≥ 72 hours) | 7.80 | 1.52-142.95 | 0.0096 | 5.83 | 1.02-111.32 | 0.047 |
| ICU at the time of TDM | 0.69 | 0.25-2.01 | 0.49 | 0.37 | 0.1-1.32 | 0.12 |
| Cefepime trough (per 1 mcg/mL increase) | 1.02 | 1.01-1.05 | 0.0016 | 1.02 | 1.00-1.04 | 0.06 |
| TDM = therapeutic drug monitoring |  |  |  |  |  |  |

Digital video EEG was recorded at the patient’s bedside in a total of 31 patients. Twelve of these patients had CIN. Most common EEG findings in patients with CIN included moderate generalized slowing of the background rhythm (11/12), generalized periodic discharges (GPDs) with triphasic morphology (9/12), rhythmic generalizing slowing not meeting criteria for GRDA (7/12) and multifocal sharp waves (6/12) (Figure 2). Non-convulsive status epilepticus (NCSE) and lateralized periodic discharges (LPDs) were found in one patient each. When looking at the combination of patterns, GPDs plus multifocal sharps/ spike and waves were the abnormalities more frequently seen together (6/12).

**Figure 2.**
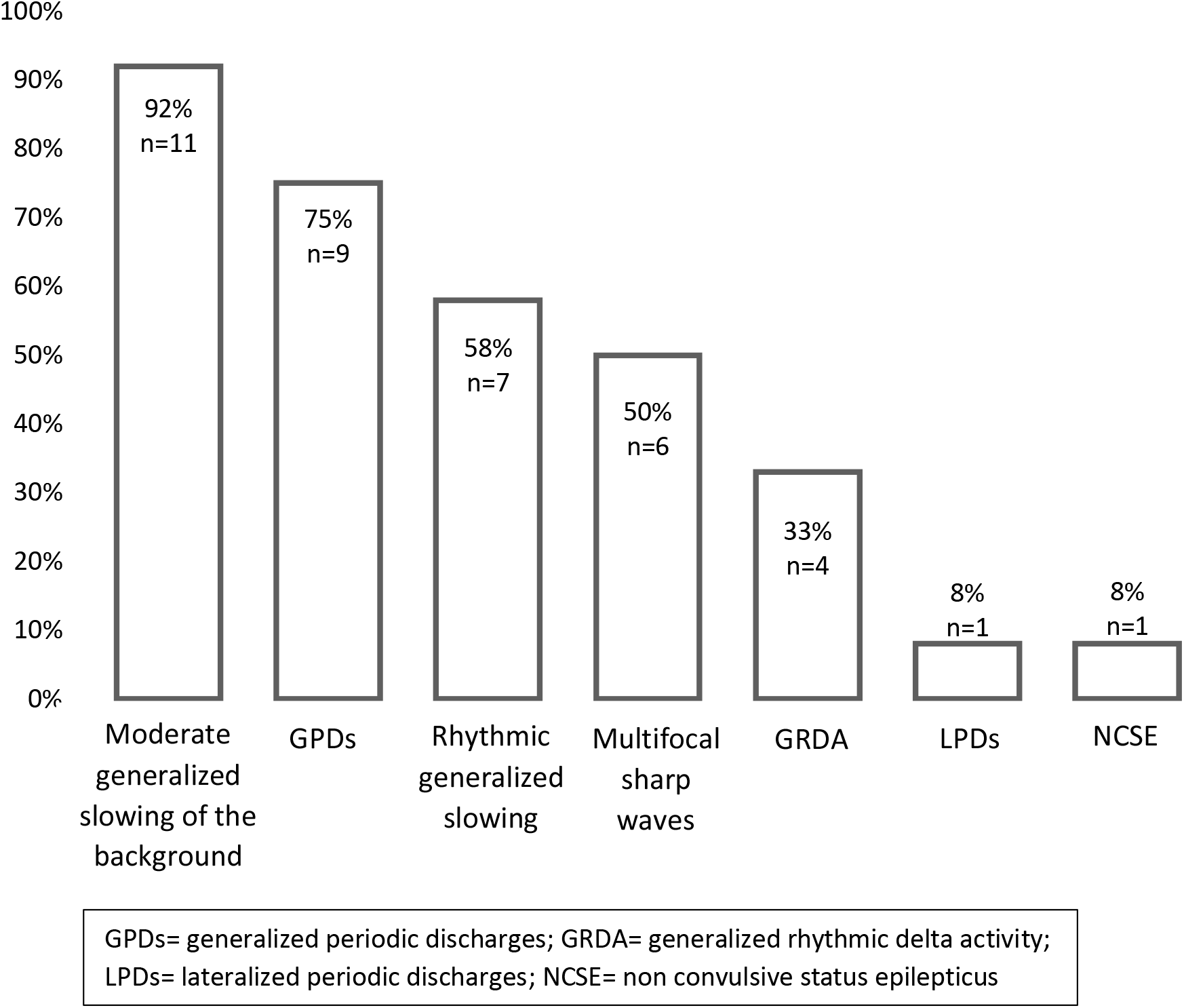
EEG Findings in Patients with CIN

## Discussion

Cefepime is a broad-spectrum antimicrobial commonly employed to combat nosocomial infections. Cefepime penetrates well into the cerebrospinal fluid (CSF) with at least 10% of drug crossing the blood brain barrier (BBB).^13^ Since cefepime is predominantly renally eliminated, a reduction in renal function increases the half-life of the drug and reduces the clearance from the body.^14^ Patients in the ICU are at a greater risk of CIN for a number of reasons such as renal insufficiency, disruptions in the BBB, and systemic inflammation resulting in increased penetration of drug into the CSF.^1^ The incidence of CIN in our study was low 13% (n=18), despite evaluation in a high-risk group (trough > 25 mg/L and ~70% in the ICU at the time of TDM). Similar to our study, Huwyler and colleagues reported neurological events in 11% of hospitalized patients receiving cefepime.^15^ Boschung-Pasquier and colleagues on the other hand reported higher rates of 23% in an inpatient population.^3^ The variability seen in the incidence of CIN rates among studies is likely due to differences in the criteria utilized to define a neurotoxic case. That said, we attribute the lower incidence of CIN in our cohort to the use of stringent criteria for identification of neurotoxicity cases.

The higher the cefepime steady state concentration, the greater the likelihood of CIN. Two previously conducted studies predicted a probability of cefepime neurotoxicity of ~ 50% if a trough of 22 mg/L was accepted as the toxicodynamic threshold.^7,8^ More recent data demonstrated cefepime trough concentrations ≥38.1 mcg/ml more frequently in those patients with presumed cefepime neurotoxicity.^3^ In sharp contrast to other published reports, the mean cefepime trough in our neurotoxicity group was much higher than reported in other studies (74.2 mg/L ± 41.1). A potential explanation for this observation is the timing of cefepime concentrations. The average time from the start of cefepime therapy to the attainment of concentrations was 6.5 days. The prolonged time to sampling likely contributed to drug accumulation and elevated concentrations particularly because most patients with CIN had diminished renal function. A noteworthy finding in this study is the association between delayed cefepime therapeutic drug monitoring (≥ 72 hours) and increased risk of CIN. There was an eight-fold increased risk of CIN when cefepime TDM was performed greater than 72 hours from the start of therapy. While beta-lactams are generally considered safe, this finding highlights the need for vigilance and early therapeutic drug monitoring to avoid overexposure of drug and minimize adverse effects. Based on the package insert, dose adjustment for cefepime is required when the CrCl is ≤ 60 ml/min.^14^ Despite having reduced renal function, the mean daily cefepime dose in the CIN group was ~4g/day, which is typically a high-dose strategy (adjusted for renal function) reserved for febrile neutropenia. The aggressive dosing utilized may indicate the growing concerns with bacterial resistance and the uncertainty in obtaining adequate drug exposure in patients with dynamic PK such as those who are critically ill.

In this study, similar to trends observed by Boschung-Pasquier et al, the in-hospital mortality rate was two-fold greater in the CIN group compared to the non-NT group.^3^ It is unknown how CIN mediates an increased risk of death. One possibility is the presence of chronic kidney disease or renal insufficiency which is an established risk factor for increased cefepime exposure. Chronic kidney disease has been associated with increased mortality due to the presence of other coexisting conditions such as cardiovascular disease and diabetes.^16^

Most of the patients in our study with CIN had an EEG performed (n=12/18, 67%) within 48 hours of the onset of symptoms and measurement of cefepime TDM. The EEG findings in this cohort are partially consistent what has been reported in the literature. The most common EEG abnormalities encountered in patients with cefepime-induced encephalopathy are generalized slowing of the background rhythm, triphasic waves and GPDs.^1,2,17^ We used the term sharp waves with triphasic morphology rather than triphasic waves as the use of the latter term is no longer recommended by the ACNS 2012 nomenclature guidelines.^11^ Other studies have encountered a higher incidence of NCSE, a difference that is also likely due to the criteria used to classify EEG abnormalities.

We recognize that there are limitations to this study. First, the retrospective study design means that causality of neurotoxicity due to cefepime cannot be fully established. The use of stringent criteria as well as review of each CIN case by three clinicians minimizes but does not eliminate the risk of misclassification. Second, the inclusion of patients who received at least 5 days of cefepime therapy means that early CIN may be under recognized in our population. The rationale for selection of this criteria was due to a previous reports of cefepime neurotoxicity in patients who received a median duration of cefepime therapy of 4 or 5 days.^1,2^ In a recent study, the median time from first cefepime dose to symptom presentation was 2 days (range 1-14 days), and this could be as short as 1 day in patients with renal insufficiency.^3,18^ Finally, we measured total cefepime plasma concentrations. We acknowledge that antibiotic protein binding in critically ill patients can be highly variable due to numerous factors such as hypoalbuminemia, renal, and hepatic insufficiency.^19^ Al-Shaer et al evaluated the total and unbound fraction of cefepime concentrations in 36 patients. Remarkably, this study revealed median fraction unbound of cefepime of 48% (range, 39%-71%).^20^ The low incidence of neurotoxicity in this study may be a reflection of this high variability in unbound drug and further emphasizes the need to measure free concentrations.

## Conclusions

Beta-lactams remain the cornerstone in the treatment of bacterial infections. Aggressive dosing strategies for beta-lactams are being utilized due to growing concerns with bacterial resistance. As cefepime doses escalate, so does the potential risk of CIN, particularly in those with renal dysfunction. In this study, we found a high rate of CIN utilizing a stringent definition for CIN. Early TDM is a valuable tool which can aid in optimizing dosing and minimizing toxicity. Further studies evaluating the pharmacodynamic relationship between cefepime concentration and toxicity are required to determine the drivers of neurotoxicity. Given the frequency of use of cefepime in the clinical setting, careful consideration of its propensity to cause neurotoxicity should be assessed prior to prescribing.

## Acknowledgments

Kenneth P. Klinker is a current employee of Merck & Co., Inc., Kenilworth, NJ. At the time of this research he was employed by the University of Florida. No funding or support was provided from Merck for this manuscript.

